# Sensory processing and participation in childhood occupation in Autism and Typically developing children - A cross sectional case control

**DOI:** 10.1101/2020.07.22.215574

**Authors:** Siew Yim Loh, Su Im Ee, Mary J. Marret, Karuthan Chinna

**Author notes:** **Corresponding author**: (SY Loh). These authors contributed equally to this work.

## Abstract

**Background:** Sensory processing difficulties and participation in childhood occupation in children impact their development, but the association among Malaysian children is unknown. The aim of this study is to provide empirical evidences on sensory processing and participation in childhood occupation, in children with autism and compare them with typical children without autism /’typical’.

**Method:** Two groups of participants (parents of children with autism and parents of ‘normal’ children were recruited from 5 hospitals, and from tuition/care centre/school respectively. Children with autism, age 6 to 10 years were matched (age/gender) with ‘typical’ children. The Participation of childhood occupation (PICO) and Sensory Processing (SSP) measures were used. Data were analysed descriptively for patterns, and Chi-square cross tabs used to compare sensory processing and participation (categorical variables) between the two groups.

**Results:** 186 parents (93 children with autism and 93 typically developing children) participated. In the autism group, 77.4 percent (n=72) were males, or 4:1 male to female ratio. Children with autism compared with typical group experienced- a) higher sensory processing difficulties and b) less participation in childhood occupation (except basic activities like eating and sleep). Sensory processing difficulties in the autism children is lower compared to developed countries, but, the prevalence of sensory processing difficulties in the ‘typical’ children (21.5 percent) was higher than data from USA and Israel (9-15%). There were significant differences in sensory processing difficulties between the two group (p<0.05), except for movement sensitivity (p=0.28). Auditory filtering section were most affected in children with autism.

**Conclusion:** Differences were found in the sensory processing difficulties (especially auditory filtering) and lower participation in autism group compared to ‘normal’ group. A higher percentage of sensory processing difficulties was also found in the ‘normal/ typically developing children’, which may be attributed to cultural or geographical factors (living in high rise flats with less playing space). More studies are needed comparing rural and urban children.

## Background

Globally, an estimate of one percent of the population is affected by Autism Spectrum Disorders and the incidences of it have risen steeply over the past decades [**1–4**]. Its prevalence is around 1.9 in every 10, 000 children before 1980, and 14.8 in every 10, 000 in 2007 [**5**], and 2.8 to 2.95 in every 10,000 in china [**6**]. Sensory Processing Difficulties (SPD) is a common sequela in children with autism. [**7–9**] SPD refers to a set of impairments where sensory information is not adequately processed, giving rise to functional difficulties in many aspects of activity participation of children with autism. In USA and Australia, there are between 69 to 95 percent of children with autism displayed symptoms of sensory processing difficulties. [**10–13**] Sensory Processing Difficulties (SPD) has debilitating consequences in children with autism as it affects activity participations such as daily living skills, academic performance [**14–15**], challenging-distressing behavior problems [**16**], and ultimately reduce the quality of life of both the child and the family [**17**]. Additionally, there is financial burdens from trying various therapies to enable these children to overcome those challenges [**18–20**], resulting in mothers of children with autism reporting more stress and depression than mothers of children without autism [**21**].

However, as there is no studies conducted on Malaysian children, this study aim to- i) examine the psychometric properties of the Malay version Short Sensory Profile and Malay version Participation in Childhood Occupation Questionnaire 2^nd^ edition (both tools will be translated and culturally adapted first); and ii) to examine sensory processing difficulties and participation in childhood occupation among Malaysian children with autism compared to typically developing Malaysian children.

The conceptual framework for the study is presented in **Figure 1**. Components of sensory processing (independent variables) is hypothesized to influence the children’s participation, with subsequent impact on their Quality of life. The ability to participate successfully in daily activities is a positive outcome of sensory integration [**17,22,23**] where any limitations in any of the component of sensory processing can affect the level of difficulty experienced when performing any activities, the frequency of performance and/or the enjoyment of activity, with impact on the quality of life [**23**].

**Figure 1.**
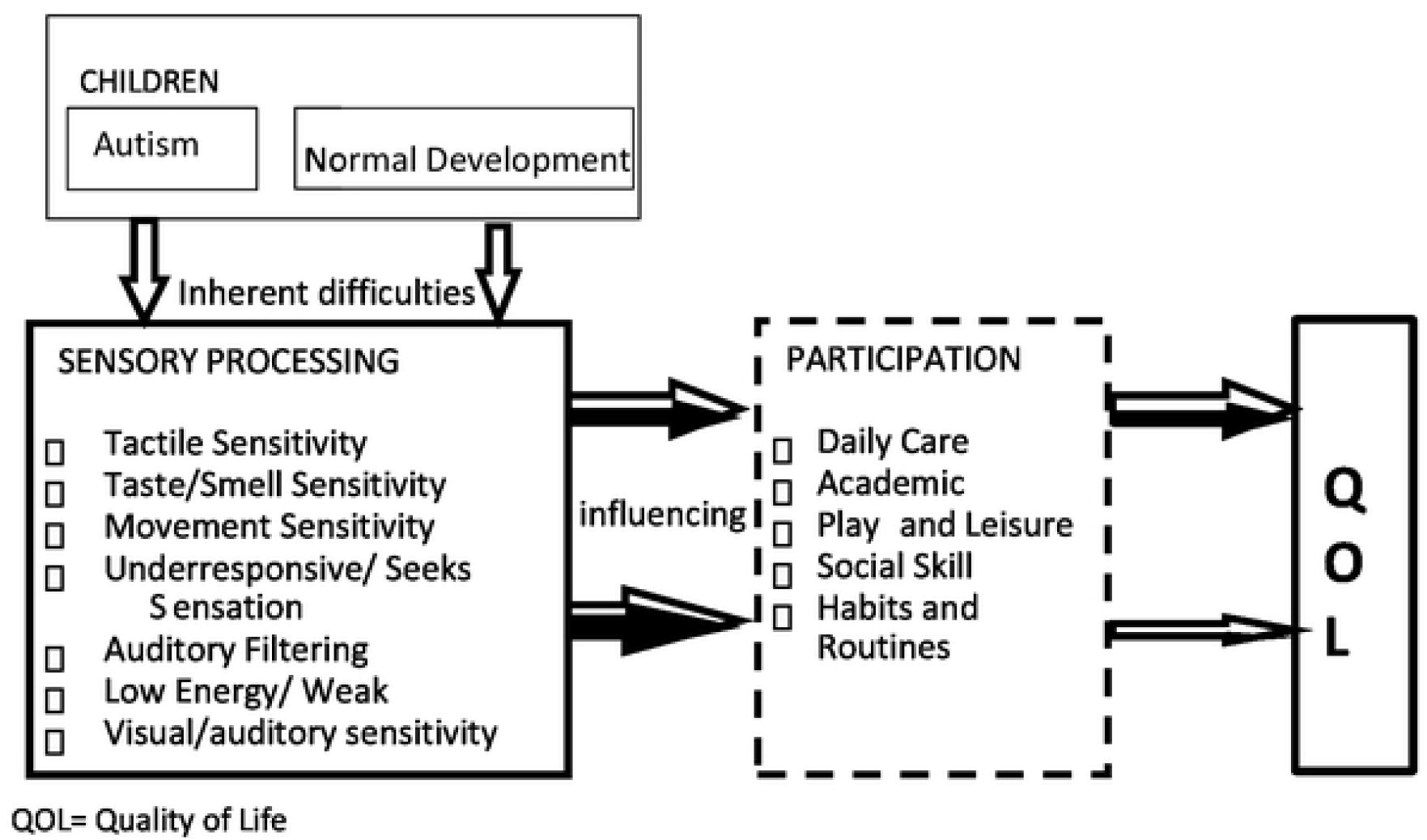
The conceptual framework with components of sensory processing (independent variables) influences activity participation of children.

## Method

This is a two phased, cross sectional study which involved translation-validating and piloting of the tool (ie phase 1 reported elsewhere), and phase two study (case control cross sectional design). The objectives of this phase two study include -i.) to examine ‘sensory processing’ patterns, the ‘participation’ patterns among Malaysian children with autism. ii) to compare the sensory processing patterns, participation patterns between children with autism and that of the typically developing children, and iii) to examine the relationship between the sensory processing pattern and the participation in activities in a cohort of children with autism. Ethical approval was obtained from the 1) Medical Ethics Committee of University Malaya Medical Centre and Research (962.25), ii) Ministry of Education (KP-BPPDP-603/5/JLD.10) to access and recruit the parents of ‘typical’ children attending schools, and iii) from the Ministry of Health (NMRR-12-1214-14435)/ MREC of hospitals to access children with autism. The Directors of five hospitals (Kuala Lumpur, Selayang, Serdang and Tengku Ampuan Rahimah Klang Hospitals).

### Participants

#### Participants

Purposive sampling was used in the data collection of children with autism, and purposive, snowball sampling was used in the data collection of typically developing children. The parents of children with autism (case group), and parents of age/gender matched ‘typical’ children (control group) were recruited. All participants were Malaysian and had children between the ages of 6 years and 10 years, matched for age and gender. Parents of children with autism were recruited from the Occupational therapy departments at five hospitals (University Malaya Medical Centre, Kuala Lumpur Hospital, Serdang Hospital, Tengku Ampuan Rahimah Klang Hospital, Selayang Hospital). Parents for the control group were recruited from tuition /after-school care centres, and school.

The criteria for children with autism is a diagnostic criteria of either the ICD 10 [**24**] or DSM-IV-TR [**25**], 6 to 10 years old, and their parents must be able to speak, read and understand Malay. The control group represent the normative sample with inclusion criteria of i) never been diagnosed a medical condition that might compromise development (e.g., attention deficit hyperactivity disorder, down syndrome, cerebral palsy, significant prematurity at 28 weeks gestation or less, very low birth weight 2000 gram or below), 6 to 10 years old and, have no sibling/s with an autism spectrum disorder that might compromise development, and their parents can speak, read and understand Malay

#### Sample Size

G* Power calculation [**26**] was performed based on the differences between the two groups (case, control) at p<0.05, 95% powers and Cohen’s d of 1.08 [**27**], on the three scales: (1) level of activity performance, (2) level of enjoyment of the activity, and (3) frequency of performance of the activity. Based on the previous means (+SD) for both groups were 125.3 (16.94) and 148.5 (10.04) in the level of activity performance, 105.1 (26.27) and 127.2 (12.11) in the level of enjoyment of the activity and for the frequency of performance 59.30 (13.95) and 67.60 (12.5) respectively. Thus, a minimum of 21 participants was required per study group to detect a difference of the defined magnitude for the scale with the highest SD (Level of Enjoyment). The second calculation was performed based on the correlation between sensory processing pattern and participation in childhood occupation for children with autism. In this calculation, conventional effect size was used since there was no previous study that examines this specific association for autism children. Assuming effect size 0.3 and 80 percent power the sample size required for this study was 84 participants with 10 percent attrition, the required sample size for children autism was 92 participants.

### Data collection

Permission to use and translate the SSP [**28**] into Malay was granted from Pearson Company [**29**]. Permission for use and translation of the PICO-Q was granted by its author [**30**]. Parents were provided with a written information sheet about the study, and asked to sign a consent form. They were assured of confidentiality, anonymity and freedom to withdraw at any time from the study with no prejudice to any future medical care.

### Data Analysis

Data analysis using SPSS Version 21, with descriptive statistics (means, frequencies and percentages) were used to analyse the demographic and patterns. Chi-square and Spearman’s correlations were used to examine significant differences between groups. Data were cleaned for outliers, and normality tests indicated a not-normally distributed group. Chi-square cross tabs were used to compare sensory processing and participation between children with autism and typically developing children with categorical variables. Spearman’s Rank-Order correlation, which is a correlation of non-parametric data, was used to examine the association between sensory processing and participation of children with autism.

### Measures

#### Demographic

A demographic and child profile questionnaire was used to gather data about the participants (children and parents of both group).

#### Short Sensory Profile (SSP)

The short sensory profile is a 38-item caregiver report to assess children’s responses to sensory events in daily life, the short version of the original Sensory Profile [31], to measure the sensory processing difficulties in children (3-10 years) with and without disabilities. It has seven sub-scales: Tactile sensitivity, Taste/Smell Sensitivity, Movement Sensitivity, Under-responsive/Seeks Sensation, Auditory Filtering, Low Energy/weak, and Visual/Auditory Sensitivity, and responses are on a 5-point Likert scale (frequently=1 to Never=5), with higher scores indicating more impairments.

The total score is calculated by the sum from the sections score. Interpretation of the total score is as follows: Definite difference indicates sensory processing difficulties, probable difference indicates questionable areas of sensory processing abilities, and typical performance indicates typical sensory processing abilities. The internal reliability (for the total and section scores) of SSP is of Cronbach’s alpha ranging from α =0.70 to 0.90. The internal validity correlations for the total and section ranged from 0.25 to 0.76. The findings from the short SSP are similar to the long version of sensory profile (Bar-Shalita, 2009). SSP has been translated into Spanish [**32**], Hebrew [**33**], Tamil [**34**] and Malay [**35**].

#### Participation in childhood Occupation (PICO-Q 2^nd^ edition)

The **Participation in Childhood Occupation [30]** is a 30-item caregiver questionnaire with five areas of functional activities: personal activities of daily living (13 items), academic activities (5 items), play and leisure (4 items), social skills (4 items), habits and routines (2 items). In addition, two questions measure the general level of participation in activities the child must perform (1 item) and activities the child chooses to perform (1 item). The 30-item PICO-Q 2^nd^ edition has additional 8 items from the initial 22-item PICO-Q, and measures five areas of functional activities (instead of four) with the additional category of social skills. Every item describes an activity that is scored against three different scales: i) level of activity performance, ii) level of enjoyment of the activity, and iii) frequency of performance of the activity. The three scales provide a score for each individual’s performance area with 15 sub-scores for the PICO-Q 2^nd^ edition.

For each item from each the three scales, parents are asked to rate using a 5-point Likert scale for level of activity performance and for level of enjoyment of the activity; while a 4-point Likert scale was used for frequency of performance of activity. The PICO-Q was tested with children with and without sensory modulation disorder and has good internal reliability with Cronbach’s alpha (alpha= 0.86 to 0.89) and good test–retest reliability (r= 0.69 and 0.86. The tool has been translated to Malay [**36**].

## RESULT

A total of 186 parents (from 93 children with autism and 93 typically developing children-age and gender matched) participated. Most of the children with autism in this study were male 77.4 percent (n=72) -ie. an approximate ratio of 4:1 male to female. **(see Table 1)**

With sensory processing, a high percentage of children with autism (68.8%) compared to typically developing children (21.5%) experienced difficulties, with total score and all sub-section scores across the SSP-M [**35**] measures were significantly different between the two groups. Children with autism experienced significantly greater sensory-processing difficulties, but the most difficult categories were *‘ Underresponsive/ Seeks-sensation’* (77.4%) and *‘Auditory-filtering’* (61.3%) while least difficult was movement sensitivity (33.3%). However, this study also found ‘sensory-processing’ difficulties for typical children were at 21.5 percent were rather high. The differences in the total SSP-M score and sub-section scores between Malaysian children with autism and typically developing children group were significant for all, except for movement sensitivity (p=0.280). For level of childhood participation in terms of i) difficulties, ii) frequency and iii) enjoyment, we found that children with autism have difficulty with social-integration; while there was a low frequency of participation in meal preparation (higher cognitive tasks), and high frequency of participation in watching videos/DVD, eating and drinking. With **enjoyment** in participation, children with autism did not enjoy social-interaction and but there was high enjoyment in solitary activities or daily routines such as watching videos/DVD on the computer or television, eating and drinking and meal time. **(see Table 2)**

With childhood participation, the **difficulties** in participation for most children with autism were in social-integration participation and participations related to homework, organizing the study environment, food preparation. There were lower frequencies in participation in food preparation, extra-curricular activities outside of the school, playing with friends, and integration in social situations, but higher frequency in solitary tasks like watching video/television. Children with autism had higher enjoyment in solitary and basic activity like video watching and meal time. children with autism have greater difficulties, lower frequencies, and less enjoyments (compared to typically developing children) across many activities. A significant correlation was found between sensory-processing (SSP-M total scores) and participation (total scores of each section in PICO2-M). More difficulties in sensory processing correlates with more difficulties in participation. auditory filtering is the only section in SSP-M that significantly correlates with all three sections of PICO2-M; difficulty in participation (r=0.36, p=<0.01), frequency of participation (r=0.22, p=<0.05) and enjoyment in participation (r=0.27, p=<0.01). **(see Table 3)**

## DISCUSSION

This is the first study in Malaysia to compare sensory processing and participation in children with autism with age/gender matched typical children. There were more males than female in the autism group, which is consistent with published gender ratios of 4 – 5 :1 [37]. Typically developing childen were age/gender matched to decrease confounding factors as patterns of participation in activity vary with age and gender [38].

### 1) Sensory processing – patterns in both group and comparisons

The Patterns of sensory processing in children with autism showed that *‘ Underresponsive/ Seeks-sensation’* (77.4%) and *‘Auditory-filtering’* (61.3%) categories were most difficult, while movement sensitivity (33.3 %) was the least difficult. Some studies are in support of this finding [**12,15**]. This finding that the sensory processing pattern among Malaysian children with autism is similar to what has been reported in western countries, suggests that culture does not affect the pattern of sensory processing in children with autism. Compared to typically-developing children, children with autism experienced greater sensory processing difficulties, in line with observations elsewhere [**39,40**]. The percentage of their sensory processing difficulties in this study is slightly lower compared to studies in the USA and Australia [**10,12,15**]. Studies in Asia do not provide prevalence or frequency of sensory processing difficulties in their samples to enable comparison [34, 41,42].

The Patterns of sensory processing in ‘typical’ children showed greater ‘sensory-processing’ difficulty at 21.5percent. However, most studies from USA reported a lower range of 5 -to- 19.9 percent [43, 44) and about 15 percent in Israel [33]. These differences may be due to cultural issues [33], believed to contribute towards the prevalence, risk and protective factors in children with disorders [45]. In addition, our study was conducted in an urban setting where a large proportion of families live in apartments, with limited opportunities for children to play outside and explore the environment. This may account for the higher percentage of typically developing children who had scores indicating sensory processing difficulties, and a useful reference for larger, future studies. With total scores and all sub-section scores of SSP-M between groups: the greatest disparity between both groups is in the auditory filtering section where children with autism were more affected. With the under-responsive/ seeks sensation category, both groups experienced high difficulties.

### 2) Patterns of participation among children with autism

This study examines the overall level of participation in terms of i) difficulties, ii) frequency and iii) enjoyment in participation in childhood occupation.

i. With **difficulties** in participation, this study found that most children with autism have difficulty with social-integration, in line with evidence that children with autism have the most difficulty in social participation [46,47]. Other difficulties in participation are related to homework, organizing the study environment, food preparation -all of which require higher cognitive abilities. Jasmin et al., (2009) [48] found that children with autism experienced severe difficulties in self-care activities, in contrast with findings by Anderson, Jablonski, Thomeer, & Knapp (2007)[49]. We found fewer difficulties with sleep, dressing, meal time and toileting too. We also found less than half of the children with autism had difficulty in watching videos/DVD on the computer or television. This is consistent with a study by Orsmond et al., (2004)[46], which indicated these video activities are prevalent in children with autism, possible because they are largely solitary activities [38,51].
ii. With **frequency** of participation, children with autism had low frequencies of participation in activities they experience difficulties, such as food preparation, extra-curricular activities outside of the school, playing with friends, and integration in social situations. Conversely there was a high frequency of participation in watching videos/DVD, eating and drinking, meal times and dressing. However, factors such as parental-involvement and access to opportunities can increase frequencies, whereby, difficult areas like socialisation that are prearranged by professionals or family members [46] can improved, even with cognitive inflexibility (likely in children with autism). Van Eylen et al., (2011) [52] suggest that proper activity instructions and the amount of disengagement may help improve participation.
iii. With **enjoyment** in participation, children with autism did not enjoy social-interaction and but there was high enjoyment in solitary activities or daily routines such as watching videos/DVD on the computer or television, eating and drinking and meal time. However. there are other confounding factors, such as family, social, sensory-behavioural challenges and communication /interpersonal relationships problems [53], which were not explored in this study.

### Participation in childhood occupation between children with and without autism

With difficulties in participation across all of the activities measured with PICO2-M, children with autism have greater difficulties, compared to typically developing children and this finding is similar to other research [54–56].

With differences in **frequency** of participation, children with autism had significantly less frequency in participation in many activities compared to typically developing children, and this finding is consistent in many other studies comparing normal children with autism and other developmental disabilities [51, 54–56]. However, there were no significant difference in the frequency of basic survival such as eating and drinking, meal time and sleep, and also in solitary tasks like watching videos/DVD’s between the two groups. Both groups in this study were reportedly flexible with unexpected changes, but as discussed, instructions given and the amount of disengagement is influential [52].

i. With differences in **enjoyment** in participation, this study found that recreational activities were enjoyed by both children with autism (87.1%) and typically developing children (93.5%), although there was no significant difference between the two groups. Some studies have found no significant difference in enjoyment of recreational activities between children with autism and typically developing children[55], while others have found that children with autism have significantly less enjoyment in participating in recreational activities [57 – 58].

#### Association between sensory processing and participation in childhood occupation

A significant correlation was found between sensory-processing (SSP-M total scores) and participation (total scores of each section in PICO2-M). More difficulties in sensory processing correlates with more difficulties in participation. There is also a significant positive but weak correlation between the enjoyment in participation section in PICO2-M (the higher score the more the child enjoys in participation) and SSP-M total scores (the higher the better sensory processing ability).

Results from this study showed that, auditory filtering is the only section in SSP-M that significantly correlates with all three sections of PICO2-M; difficulty in participation (r=0.36, p=<0.01), frequency of participation (r=0.22, p=<0.05) and enjoyment in participation(r=0.27, p=<0.01). Auditory filtering difficulties associated with problems in learning and attention and thus may cause poor academic performance in children with Autism [14].

Low energy and tactile sensitivity were significantly correlated with difficulties and enjoyment in participation but not with frequency of participation. Previous studies also suggested that tactile sensitivity can affect daily activities [59] Jasmin et al., 2009 [48] and academic and social activities [7]. Therefore, sensory processing difficulties is one of the important factors that may contribute to difficulties and enjoyment in participation.

Occupational therapist helps the children with autism to engage in activities participation by reducing the barriers, without giving too much stress or burden to the family. Studies on correlation between sensory processing difficulty and patterns of participation in childhood occupation are important and coincide with the concept of the World Health Organization [60] which emphasises the importance of the relationship between disability, performance and participation in daily living activities, outlined in this study’s conceptual framework.

#### Strength & Limitation

The study is a cross sectional design study with an urban sample, and a reliance on report from parents. Therefore, the findings may have limitation for generalisation. Future studies should include, parent reports, with reports from teachers, and clinical observations for a more balanced profile of the child. Future studies should also sample from both urban and rural settings, with more representation from a multi-ethnic Malaysian sample.

## Conclusion

This is the first case control study conducted in Malaysia, where significant differences in pattern of sensory processing and, pattern of participation in childhood occupation were found between children with autism and ‘normal’ children. A high percentage of children with autism (68.8%) experienced sensory processing difficulties which is also reported in 21.5 percent of typically developing children. This calls for clinician to assess for these deficits which affect daily performance. Children with autism showed difficulties in seeking sensation, auditory filtering and social situations (eg playing with friends), being responsible for homework, organizing the study environment and preparing simple food. Activities which most children with autism experience less difficulty are those with a fixed sequence of steps and has less demand on cognition.

Auditory filtering (of sensory processing) is the only section in SSP-M that shows correlation with all three sections of PICO (participation), and the correlation is highest with the difficulty in participation total score. A moderate to weak but positive correlation between sensory processing (SSP-M total score) and childhood participation (PICO2-M sections total score) suggests that addressing sensory processing alone will not necessarily result in an improvement in participation in childhood occupation. More studies are needed to examine all related factors that may contribute to limitation in participation.

